# Constraints and limitations on the transcriptional response downstream of the Bicoid morphogen gradient

**DOI:** 10.1101/728840

**Authors:** Huy Tran, Aleksandra M. Walczak, Nathalie Dostatni

## Abstract

The regulation of the *hunchback* promoter expression by the maternal Bicoid gradient has been studied as a model system in development for many years. Yet, at the level of quantitative agreement between data and theoretical models, even the first step of this regulation, transcription, continues to be challenging. This situation is slowly progressing, thanks to quantitative live-imaging techniques coupled to advanced statistical data analysis and modelling. Here we outline the current state of our knowledge of this apparently “simple” step, highlighting the newly appreciated role of bursty transcription dynamics and its regulation.

## 1. Introduction

The development of properly proportioned individuals relies on cells precisely reading out positional information and committing to a specific cell fate. In a developing organisms, non-uniformly distributed morphogens gradients provide such positional information to each cell by turning on the expression of specific target genes in a concentration-dependent manner (Karaiskos et al., 2017; X. Liu et al., 2009; Petkova, Bialek, Wieschaus, & Gregor, 2019). Although these gradients are known to be essential since decades in many developmental systems, in most cases we do not know how they are established or how they activate target gene expression in a dose-dependent manner. Here, we discuss recent developments in describing how a noisy transcriptional process can be tuned very rapidly into a reproducible functional developmental pattern by focusing on one well-studied morphogen-target gene pair – the Bicoid (Bcd) morphogen controlling the *hunchback* (*hb*) gap gene in early fly development.

The *hb* gap gene is involved in antero-posterior (AP) patterning of young fruit fly embryos. *hb* expression is first detected at the onset of zygotic transcription (nc 8), around one hour after fertilization. No more than 30 min later (nc 11), *hb* transcription occurs in an anterior domain with most nuclei highly expressing the *hb* gene. The border separating this expressing anterior domain from the posterior domain with mostly *hb*-silent nuclei is extremely sharp (Little, Tikhonov, & Gregor, 2013; Porcher et al., 2010) (Figure 1A). This type of expression pattern defines a step-like pattern. Following the *hb* transcription step-like pattern, the Hb protein expression quantified at nc14 is also step-like (Houchmandzadeh, Wieschaus, & Leibler, 2002) and exhibits very low variability in concentration between nuclei at the same position along the AP axis (Gregor, Tank, Wieschaus, & Bialek, 2007). This precise Hb pattern, combined with other gap gene patterns, was proposed to contain enough positional information for nuclei to predict their position in the embryo with ∼ 99% accuracy (Dubuis, Tkacik, Wieschaus, Gregor, & Bialek, 2013; Petkova et al., 2019).

**Figure 1.**
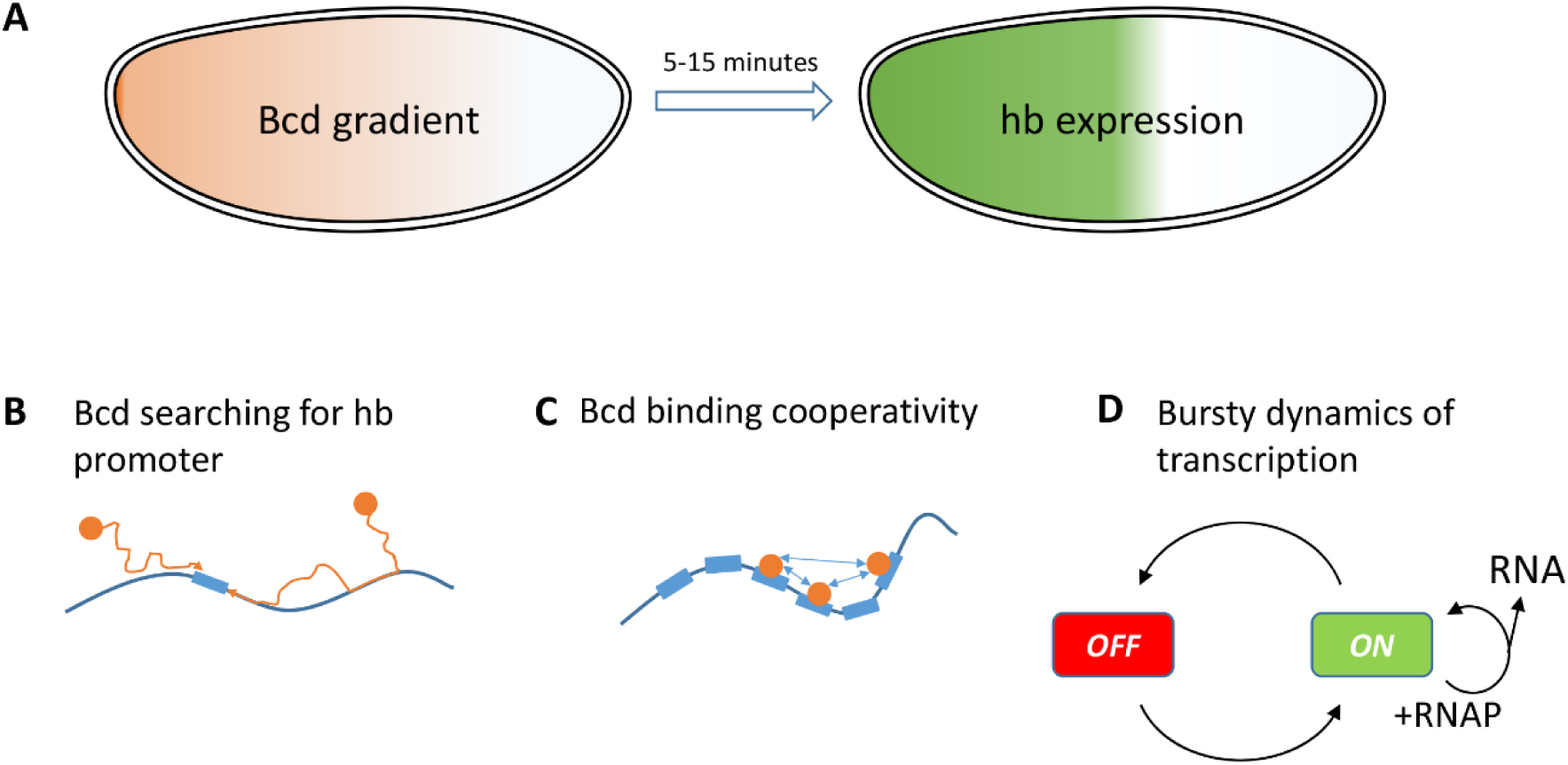
**A**) How is positional information contained in the shallow Bcd concentration gradient transformed into a robust step-like *hb* expression pattern in the 5 to 15 minutes of the nuclear cycle 11-13? **B-D)** The rate-limiting steps of *hb* expression in response to the Bcd gradient: **B:** The Bcd searches for the *hb* promoter, which can involve either 3D diffusion inside the nuclear space or 1D diffusion along the DNA. **C:** Bcd binds cooperatively to *hb* promoter, separating the embryo into a rich Bcd-bound anterior region and a no Bcd-bound posterior region. This process can involve exclusively Bcd molecules, other DNA bound transcription factors or factors facilitating chromatin accessibility. **D:** The production of *hb* RNA was shown to be bursty (Desponds et al., 2016), with relatively infrequent switching of the *hb* promoter between the ON and OFF expression states.

In young embryos, the major regulator of *hb* transcription is the homeodomain-containing Bcd (Driever & Nusslein-Volhard, 1988). Translated from maternal mRNAs anchored at the anterior pole, Bcd proteins form an AP exponential gradient and activate *hb* transcription in the anterior half of the embryo. Increasing or decreasing the amount of Bicoid in the embryo induces a posterior or anterior shift, respectively, of the *hb* step-like pattern, arguing that the expression of *hb* is dependent on Bcd concentration (Driever & Nusslein-Volhard, 1988). The discovery of Bcd’s role in *hb* transcription was exciting as it provided the first example illustrating the elegant idea of the French Flag model for morphogenesis. However, 30 years after its discovery, the mechanisms responsible for the sharpness of the *hb* step-like pattern and its reproducibility are not yet fully understood. The first quantitative studies of the Bcd gradient and Hb protein pattern reproducibility (Gregor, Wieschaus, McGregor, Bialek, & Tank, 2007; Houchmandzadeh et al., 2002) raised doubts about the pattern emerging solely from diffusive biochemical interactions between transcription factors and the gene promoter region. Gregor et al. (2007) used the Berg-Purcell scheme (Berg & Purcell, 1977), in which the concentration of a diffusing ligand is estimated by counting the number of ligand-receptor binding events within a specific time. In this simplest scheme of concentration sensing, the time required for a Bcd binding site of the *hb* promoter to read the Bcd concentration with 10% accuracy was estimated to be of the order of 2 hours (Gregor, Tank, et al., 2007), much longer than the interphase duration of nuclear cycles. These early attempts to explain the establishment of the Hb step-like pattern marked a shift in experimental studies towards more quantitative approaches. The advent of new measurement methods, based on live imaging especially of transcription dynamics (Bertrand et al., 1998), allowed for a deeper view of the processes of positional readout in young fruit fly embryos. Specifically it allowed one to focus on the outcome of the first steps of regulation, transcription at the *hb* locus, to determine how this stage of early development is controlled. Thanks to advances in imaging discussed below, the pattern of transcription from the *hb* locus is observed to be established sharply in about 3 min at nc11, despite being generated by noisy gene expression from a bursty promoter. In this review we discuss recent work and challenges for advancing our understanding of positional information propagation from the Bcd gradient to the *hb* expression pattern (Figure 1). We made the choice to only focus on the first step, the transcription process, which in itself contains several steps (Figure 1B-D), and highlight why this system remains challenging 30 years on. More details about the establishment of the Bcd gradient can be found in the chapter by Huang and Saunders in the same issue (Huang & Saunders, 2019).

## 2. The Bcd gradient

### 2.1. Positional information

The Bcd concentration gradient arises from maternal mRNA anchored at the anterior pole of the syncytial embryo (Grimm, Coppey, & Wieschaus, 2010). Large scale analysis of the Bcd gradient, through immunofluorescent staining of the endogenous protein or analysis of the fluorescent-fusion Bcd-eGFP protein (Gregor, Wieschaus, et al., 2007) (Figure 2A), indicated that the Bcd gradient is exponential with a concentration decay length around 80 to 120μm (between one-fifth and one-fourth of embryo length) (Abu-Arish, Porcher, Czerwonka, Dostatni, & Fradin, 2010; Gregor, Bialek, de Ruyter van Steveninck, Tank, & Wieschaus, 2005; Gregor, Wieschaus, et al., 2007). The concentration of this gradient is a source of positional information for each nuclei along the AP axis. All the experiments performed to measure the absolute concentration of the Bcd gradient along the AP axis used the Bcd-eGFP expressed from a transgene in the embryo which rescues the viability of the *bcd*^*E1*^ null allele. The total concentration of Bcd at the anterior pole, measured *via* fluorescent Bcd-eGFP ranges from 90 nM, measured by comparing the fluorescence inside nuclei with the fluorescence of an eGFP solution at a given concentration (Gregor, Garcia, & Little, 2014), to 140 nM (Abu-Arish et al., 2010), equivalent to respectively, 4.4 (estimated to be ∼ 700 Bcd molecules per nuclei at nc14 (Gregor, Wieschaus, et al., 2007)) to 7 molecules/*µm*^3^ where the *hb* pattern boundary is established. Even though these measurements are consistent, they were obtained with the same fluorescent Bcd-eGFP and are likely to be underestimates of the real concentration because a proportion of the eGFP might not be fluorescent. Further analysis using a Bcd fusion carrying two fluorescent domains (Durrieu et al., 2018) might help resolve the issue of absolute Bcd concentration measurements in the embryo. Similarly, taking into account the maturation time of the eGFP points to a slight overestimation of the Bcd gradient decay length of about 15% (Durrieu et al., 2018; F. Liu, Morrison, & Gregor, 2013). After corrections, the length constant of the Bcd gradient is estimated as low as one sixth of embryo length (16.5 ± 0.7 % EL).

**Figure 2.**
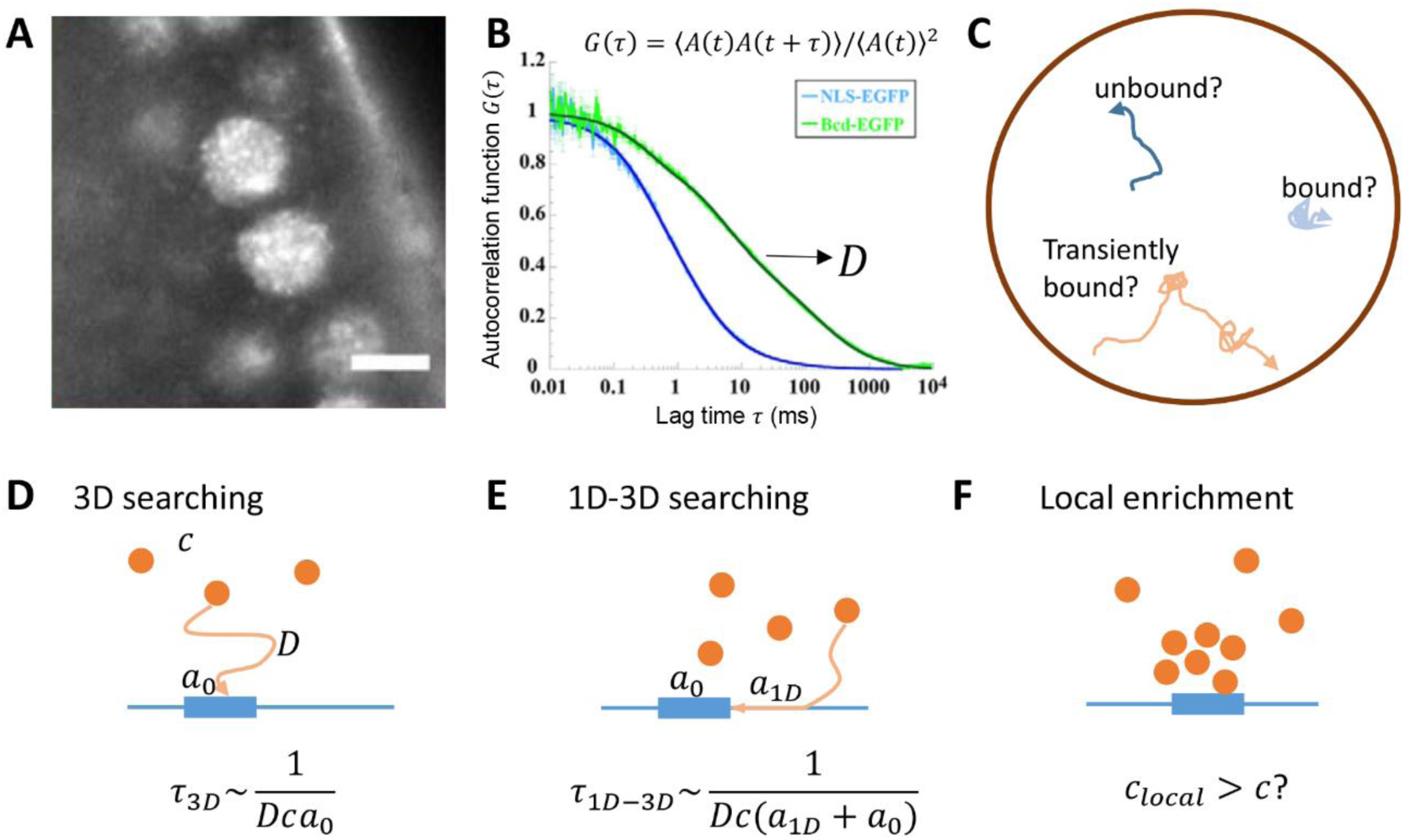
Studying Bcd motility in living fruit fly embryos. **A)** Fluorescent-tagged Bcd molecules were shown to be distributed in foci in the nuclear space of fixed embryos (H. Xu, Sepúlveda, Figard, Sokac, & Golding, 2015) and hubs of fluorescently-tagged Bcd (Mir et al., 2017, 2018). Figure reused from (Mir et al., 2017). **B-C**) Different motilities of Bcd molecules were inferred from Fluorescent Correlation Spectroscopy (FCS) analysis of the Bcd-eGFP signal in nuclei, with a Bcd population with a high (free) diffusion coefficient (**B**), Figure reused from (Porcher et al., 2010) and **C**, by single-molecule tracking approaches revealing bound, unbound and transiently bound molecules (Mir et al., 2018). **D-F)** various scenarios of Bcd searching for its binding sites in the *hb* promoter. **D**: Bcd molecules can diffuse in 3D nuclear space to search for Bcd binding sites on the target promoter. The time, ***τ***_**3*D***_, for the binding site to be found is inversely proportional to the product of *D* (coefficient of free diffusion), *c* (concentration) and *a*_0_ (size of the binding site); **E**: Bcd molecules can “hop” to non-specific locations on the DNA and slide along DNA segments in 1D to search for specific binding sites. The average sliding distance (or footprint) ***a***_**1*D***_ can be much bigger than the size of a Bcd binding site and therefore reduces the search time from the 3D case; **F**: Local enrichment of transcription factor (TF) concentration (*c*_local_>*c*) at the promoter region can also reduce the promoter search time.

### 2.2. The motility of Bcd molecules

The *hb* locus extracts positional information from the local Bcd concentration via interactions with Bcd molecules. Given the short time window for positional readout in each interphase, the Bcd search time for the *hb* promoter *τ*_*search*_ is critical in determining the limit of positional readout precision. Therefore, several studies analyzed Bcd motility, using FRAP (Gregor, Tank, et al., 2007) or FCS (Abu-Arish et al., 2010) on fluorescent Bcd-eGFP or single-molecule tracking (Figure 2A) (Drocco, Grimm, Tank, & Wieschaus, 2011; Mir, Stadler, Harrison, Darzacq, & Eisen, 2018)). In the initial FRAP experiments, Bcd motility in the cytoplasm turned out to be quite slow (∼ 0.3 µm^2^/s) (Gregor, Wieschaus, et al., 2007). FCS experiments performed both in the cytoplasm and the interphase nuclei revealed the existence of Bcd molecules with different motilities: best fitting of the data to the two-species diffusion model indicated that *i)* in the cytoplasm, 18% of the Bcd molecules are slow-moving while 82% of the Bcd molecules are fast-moving with an average diffusion coefficient of ∼ 7.4 µm^2^/s (Abu-Arish et al., 2010) and *ii)* in the nucleus, 43% of the Bcd molecuels are slow-moving (∼ 0.22 µm^2^/s) and 57% fast-moving (∼ 7.7 µm^2^/s) (Porcher et al., 2010). Fast moving Bcd molecules (∼ 4 µm^2^/s) were also observed using a photoactivable Dronpa-Bcd (Drocco et al., 2011) and the existence of at least two populations of Bcd molecules was further confirmed by high resolution single molecule imaging suggesting that in nuclei, Bcd molecules spend the same amount of time on nuclear exploration (searching for a binding target) and on binding to chromatin with surprisingly high unbinding rates, distributed with long tails (Mir et al., 2018).

### 2.3. The time it takes for Bcd to find its target sites on the *hb* promoter

Transcription factors (TFs) explore, usually by passive diffusion, the nuclear space in “search” of their DNA binding sites on the regulatory sequences of their target genes. In the case of *Drosophila* embryogenesis, the time it takes for the Bcd protein to find its target sites on the *hb* promoter is critical to explain how *hb* expression in response to Bcd can be so fast yet precise despite the limited Bcd concentration in the mid-embryo region. Therefore, many studies have used modeling to estimate the time it takes Bcd to find its target sites and proposed several strategies to shorten this search time (Abu-Arish et al., 2010; Gregor, Tank, et al., 2007; Mir et al., 2017, 2018; L. Mirny et al., 2009; Normanno, Dahan, & Darzacq, 2012).

In an initial attempt to estimate the Bcd search time for the *hb* promoter, Gregor and colleagues proposed that Bcd molecules can diffuse in 3D inside the nuclear space to come in contact with the specific Bcd binding sites on the *hb* promoter (Figure 2D). They proposed that the lapse time between each contacts was inversely proportional to the diffusion coefficient of Bcd *D*, the local Bcd concentration *c* and the size of the Bcd binding site *a*.

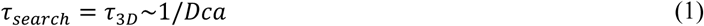

The binding site for TFs, including Bcd, are commonly 10bp long which corresponds to ∼3nm. Given this value and assuming that the searching Bcd molecules are diffusing freely (i.e. are fast-moving molecules) and that Bcd binding is diffusion limited (each collision between a Bcd molecule and a binding site results in a successful binding event), the binding site search time *τ*_3*D*_ is estimated to be on the order of 10 s. However, this is an optimistic estimate given that *a* is likely 10 fold too large since displacement by a single bp may lead to an entirely different DNA sequence that is not recognizable by the protein (Slutsky & Mirny, 2004). In this later case, *a* would be ∼ 0.3 nm and the search time for Bcd would be 100s. In addition, activation by Bcd requires in general not just the binding of one Bcd molecule to the *hb* promoter but several (Burz, Rivera-Pomar, Jäckle, & Hanes, 1998; Driever, Thoma, & Nüsslein-Volhard, 1989; Ronchi, Treisman, Dostatni, Struhl, & Desplan, 1993). Thus, clustering of binding sites in the regulatory sequences of the target genes is another factor that may lengthen the time it takes to bind enough Bcd molecules together to get a transcriptional response.

The rapid establishment of the *hb* pattern, despite a long search time in 3D, leads to hypotheses that Bcd molecules may use a combination of 1D and 3D diffusion to search for their target sites (L. Mirny et al., 2009; Slutsky & Mirny, 2004). This mode of searching was first observed in *Escherichia coli* (Elf, Li, & Xie, 2007; Hammar et al., 2012; Riggs, Bourgeois, & Cohn, 1970) for the *lacI* repressor where a combination of a 1D and 3D search was found to reduce the search time by up to 100 fold compared to a pure 3D search. In this scheme, proteins bind non-specifically to DNA and, while doing so, slide along the DNA segment in search for the specific target site (Figure 2E). Therefore, the effective size of the target site is increased by the TF’s sliding footprint along the DNA *a*_1*D*_ (up to a hundred bp, compared to the size of a binding site *a* = 10 bp). Another hypothesis is that the target locus could be located in a micro-environments with enhanced TF concentration (*c*_local_>*c*) (Figure 2F), thus speeding up the search, as proposed in the case of the Ultrabithorax protein (Tsai et al., 2017). The recent observation of Bcd concentration in dense hubs (Mir et al., 2017, 2018) opens up the possibility that micro-environments that enhance local Bcd concentration could contribute to reduce the Bcd search time for the *hb* promoter. However, one should note that these mechanisms can also introduce non-linearities into the position sensing process: Bcd hubs were found to persist even in the posterior region of low Bcd concentration, leading to a much flatter Bcd concentration profile in hubs than in the cytoplasm. In addition, Hammar *et al.* observed that bound lacI molecules may interfere with the 1D-sliding molecules when the distance between the target sites is shorter than the sliding footprint (Hammar et al., 2012). If Bcd employs this mode of searching, the very short distances between Bcd binding sites on the *hb* promoter (as short as 12 bp) may introduce negative feedback to Bcd binding, instead of the positive feedback normally linked to a sharp *hb* pattern (Ma, Yuan, Diepold, Scarborough, & Ma, 1996). It is experimentally challenging to directly observe the scheme of the Bcd target site search. Currently, it is not possible using a confocal microscope to directly identify which one of the Bcd subpopulations with characterized motilities are directly searching for target sites. Observing interactions between Bcd and its target sites is still limited to epigenomics approaches such as ChIP on fixed samples (Bradley et al., 2010; Li et al., 2011) performed on population of nuclei that have very different transcription features. Recent advances in sample preparation (Combs & Eisen, 2013; Karaiskos et al., 2017) may allow us to observe how these interactions correlate with the varying degree of inhomogeneity in Bcd motility along the embryo AP axis and help identify the target site search mode.

### 2.4. Activation of transcription by Bcd

The Bcd protein is able, on its own, to activate transcription when bound to a promoter containing as few as three of its DNA binding sites (Crauk & Dostatni, 2005; Ronchi et al., 1993). However, how this is achieved remains largely unknown. Structure-function analyses of the Bcd transcription factor indicated that it contains many redundant functional domains (Schaeffer, Janody, Loss, Desplan, & Wimmer, 1999). Besides its homeodomain which allows binding to DNA (Hanes & Brent, 1989; Trelsman, Gönczy, Vashishtha, Harris, & Desplan, 1989), the Bcd protein contains several independent activation domains which can activate transcription on their own when multimerized and fused to a Gal4 DNA binding domain in vitro (Sauer, Hansen, & Tjian, 1995) or in the early embryo (Janody, Sturny, Schaeffer, Azou, & Dostatni, 2001). These include a Glutamine-rich domain, a ST-rich domain and a C-terminal acidic domain. The Bicoid protein also contains independent inhibitory domains which reduce its activation potential (Janody et al., 2001; C. Zhao et al., 2002). Finally, activation by Bcd is enhanced by other TFs binding to the promoter. These include the maternal contribution of the Hunchback protein itself (Porcher et al., 2010; Simpson-Brose, Treisman, & Desplan, 1994) or Zelda (Mir et al., 2017; Z. Xu et al., 2014). Yet, the mechanisms underlying these essential synergistic effects are poorly understood.

## 3. *hb* transcription dynamics

### 3.1. Visualizing *hb* transcription dynamics

RNA-FISH on fixed embryos allowed for the monitoring of *hb* nascent transcript accumulation at their site of synthesis inside each nucleus of the embryo, making it an initial marker to study ongoing transcription and promoter dynamics at a given locus. This allowed subsequently for the detection of single mature mRNAs in the cytoplasm and in the nucleus (Little et al., 2013; Perry, Bothma, Luu, & Levine, 2012; Zoller, Little, & Gregor, 2018). Distributions of nascent and cytoplasmic *hb* RNA along the AP axis demonstrate sharp step-like patterns at the transcription level, with an expression boundary width varying from 10 % egg length to 8% egg length at nc11 and nc13 respectively. The observed data suggested, despite low heterogeneity at the protein level (Gregor, Tank, et al., 2007), a very noisy transcription process occurring with periods of promoter activity and inactivity (H. Xu et al., 2015).

RNA FISH requires fixation of the sample and can only provide a snap shot view of the transcription process at a given time (the time of fixation) during nuclear interphase. Following the pioneering work of R. Singer (Bertrand et al., 1998), the MS2 fluorescent RNA-tagging system has been implemented to monitor transcription dynamics in living early *Drosophila* embryo development (Garcia, Tikhonov, Lin, & Gregor, 2013; Lucas et al., 2013). The system takes advantage of strong interactions between the MCP coating proteins and its RNA stem loops from the MS2 bacteriophage. As nascent RNA containing stem loops are being transcribed, they are bound by fusion proteins MCP-GFP, making ongoing transcription at the loci visible as bright fluorescent spots under the confocal microscope (Ferraro, Lucas, et al., 2016).

Recently, the MS2 system allowed for the direct visualization in real-time of position-dependent activation of the proximal *hb* P2 promoter (700 bp) (Lucas, Tran et al., 2018): at each nuclear interphase (from nc11 to nc13), *hb* expression (detected via MCP-GFP foci) first occurs in the most anterior then proceeds progressively though very rapidly from the anterior to the boundary region (at ∼ 45% of embryo length from the anterior pole). Of note, this progressive appearance of first *hb* foci after mitosis is measured considering as an origin of time the onset of mitosis of each nucleus and is thus not a consequence of the mitotic wave. This dynamics from the anterior pole to the center of the embryo occurs even within the anterior region where Bcd is presumably saturated. These findings suggest that Bcd concentration is not only rate-limiting in the boundary region, as observed from the pattern dynamics at the steady-state, but also within the anterior region. In this region, differences in Bcd concentrations leads to differences in *hb* activation times (up to 3 minutes), which can take significant portion of the early nuclear interphase and consequently affects the total amount of transcripts produced. In addition, analysis of the MCP-GFP time trace of each individual nucleus shows variability in the transcription process even in the anterior region with high Bcd concentration (Desponds et al., 2016), indicating that Bcd is not the sole factor contributing to the noise in *hb* transcription.

### 3.2. Characterizing *hb* transcription dynamics

The fluorescent time traces acquired with the MS2 system provide an indirect observation of transcription dynamics. The signal is noisy, convoluting both experimental and intrinsic noise with the properties of the MS2 probe. To obtain a specific fluorescent signal sufficiently strong to overcome background fluorescence due to unbound MCP-GFP molecules, a long probe of 24 RNA stem loops was used (Garcia et al., 2013; Lucas et al., 2013). As the signal is only detected while the probe is being transcribed (Figure 3A and B), this introduces a significant delay (buffering time) between each instant the promoter is ON and the corresponding fluorescent detection. This buffering time of ∼1 min (Coulon et al., 2014; Fukaya, Lim, & Levine, 2017; Garcia et al., 2013) and the short length of the traces (5-15 minutes) prevent traditional analysis based on OFF time distributions or autocorrelation functions to quantify the statistics of the activation and inactivation times.

**Figure 3.**
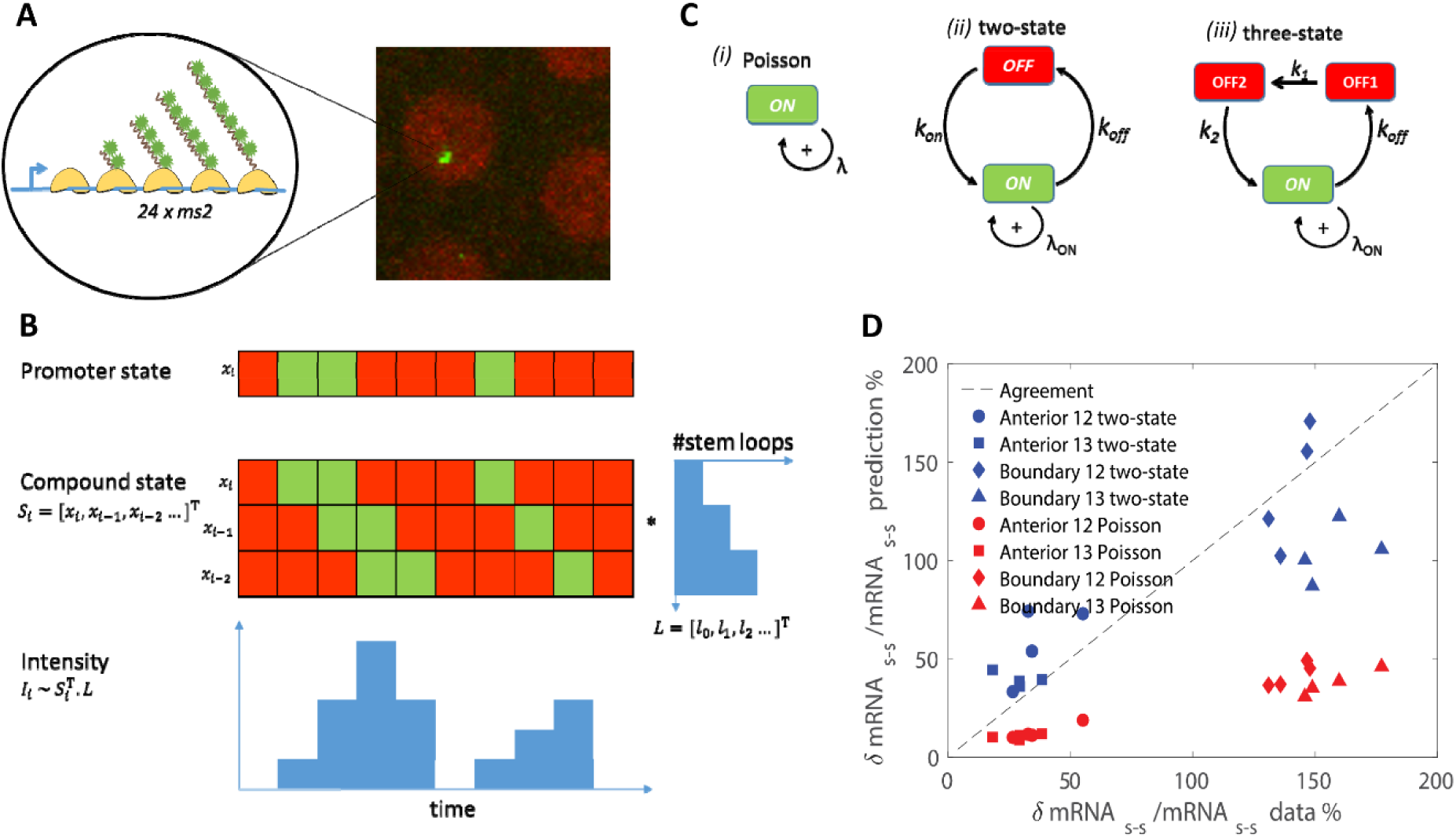
Inferring promoter dynamics from MS2 loci intensity: **A**) Visualization of active transcription loci: as RNAs containing MS2 stem loops are transcribed by RNA Polymerases (RNAP, in yellow), they are bound by fluorescent MCP-GFP molecules (in green). The succession of several RNAPs transcribing the gene allows for the accumulation of several fluorescently tagged MS2-containing RNA at the same location which become visible under the confocal microscope as green bright spots (right panel). **B**) Transformation from the promoter state dynamics to MS2 spot intensity dynamics: promoter state in discrete time ***x***_***i***_ indicates whether the promoter is ON (green) or OFF (red). This state is encoded in the compound state vector ***S***_***i***_ **= [*x***_***i***_, ***x***_***i***−**1**_, ***x***_***i***−**2**_ **…]**, which also maps the position of RNAP on the MS2 cassette. RNAP arriving at time *i* will be transcribing a nascent RNA containing ***L***_***j***_ MS2 stem loops at time ***i* + *j. L*** depends on the length and on the arrangement of the MS2 stem loops on the reporter gene. The active loci intensity ***I***_***i***_ at time ***i*** is given by the product of ***S***_***i***_ and ***L*. C**) Different models of promoter dynamics: *i)* Poisson model: random RNAP arrival and initiation of transcription; *ii)* two-state and *iii)* three-state models, where promoters switch successively between ON and OFF states. During the ON state, RNAPs arrive and initiate transcription in bursts, with maximum rates. **D**) Comparison of readout noise (***δmRNA/mRNA***) at steady state generated from Poisson (red) and bursty two-state models (blue) and data (dashed). Figure reused from (Desponds et al., 2016).

Desponds et al. developed a tailored autocorrelation analysis of the fluorescent time traces to overcome these limitations (Desponds et al., 2016). Combining this analysis with models of transcription initiation (Figure 3C) and estimates of the precision of the transcriptional readout, provided evidence for bursty transcription initiation in nuclear cycles 12-13 (Desponds et al., 2016). Namely, they find the dynamics in agreement with a telegraph model, in which the promoter switches between the ON and OFF states. Only during the ON state can RNA polymerase arrive and initiate transcription successively (Figure 3C). The switching between the ON and OFF promoter states can be driven by polymerase pausing which was proposed to prevent new initiation between transcription bursts (Shao & Zeitlinger, 2017), the binding/unbinding of other TFs required for activation, or DNA looping for distal enhancer-promoter contacts (Fukaya, Lim, & Levine, 2016). The best-fit switching period (for a full ON-OFF cycle) is in the order of ∼30 s, with the probability to be in the ON state of ∼50% at the anterior and of ∼10% at the boundary.

It should be noted that the autocorrelation function analysis alone is not able to distinguish reliably between different models for promoter activation and requires complementary information about the precision of the transcriptional readout to conclude that transcription is most likely bursty (Figure 3D). Recently, an inference method based on hidden Markov model and maximum likelihood has been developed (Corrigan, Tunnacliffe, Cannon, & Jonathan, 2016) and tailored for the MS2 system in *Drosophila* (Lammers et al., 2019). This discrete-time model employs a hidden compound state, which records the previous promoter states during the elongation time (Figure 3B). This compound state is used to map RNA Polymerase (RNAP)’s position on the reporter gene segment and calculate the active loci intensity at a given time. The rates of switching between the promoter states in each time step are fitted based on maximum likelihood. While computationally expensive, the method allows for direct model selection and shows that transcription bursts are prevalent in stripe gene expression in later stages of fly development (Berrocal, Lammers, Garcia, & Eisen, 2018; Bothma et al., 2014; Lammers et al., 2019).

### 3.3. Transcription regulation of *hb* gene by Bcd proteins

Data obtained from the MS2 system provided insights not only at the molecular level about the kinetics of the promoter behavior but also at the cellular level when considering individual nuclei along the AP axis and individual loci in each of these nuclei. In particular, it was possible to analyze the transcription dynamics of each *hb-MS2* locus at the scale of the whole embryo. This analysis indicated that depending on its position along the AP axis, each locus was able to either turn ON when positioned in the anterior or remain silent when positioned in the posterior. Surprisingly, the steep border forms in under 3 min at each nuclear interphase 11 to 13 (Lucas, Tran et al., 2018). This indicates that the system is able to measure extremely rapidly very subtle differences of Bcd concentration and produce a complete sharp border. This rapid responsiveness is fascinating because it is almost ten times faster than predicted by previous theoretical models assuming that the Bcd gradient is the only driver for the *hb* transcription process.

The steep Bcd-dependent *hb* pattern, given the smooth Bcd gradient, demonstrates a strong nonlinear regulation of the *hb* gene by Bcd. The presence of multiple Bcd binding sites on the *hb* promoter (Driever & Nusslein-Volhard, 1988) suggests that such strong nonlinearities can be achieved by high cooperativity of Bcd binding to the *hb* promoter site. Cooperative binding of Bcd to multimerized binding sites was observed *in vitro* (Burz et al., 1998; Ma et al., 1996) but remains too weak to account for the extremely steep Bcd-dependent *hb* pattern observed *in vivo*. Synthetic reporters with only Bcd binding sites are weakly expressed in very anterior domains which harbor, however, remarkably steep posterior boundaries (Crauk & Dostatni 2005; Ronchi et al. 1993). This suggests that Bcd and Bcd binding sites are sufficient to generate a steep posterior border and models of regulation by Bcd binding/unbinding can help understand how this is achieved.

Using powerful optogenetics to manipulate Bcd activity in real time, Huang and colleagues demonstrated that the full functional Bcd is critical in the early nuclear cycles (nc11 to 13) for the development of the embryo’s mesothorax, where the *hb* boundary is established (Huang, Amourda, Zhang, Tolwinski, & Saunders, 2017). During this time window, light-induced conformational changes of the Bcd molecules, which leads to their functional inactivation, significantly reduces transcription at the *hb* promoter, as shown using the MCP-GFP system. This study links the transcriptional activity of TF bound to their target sites with the temporal dynamics of transcription and provide an unprecedented time resolution of the TF-dependent transcription process. A general model of transcription regulation via binding/unbinding of TF to the binding sites on the target promoter (Figure 4A), demonstrates that the pattern’s degree of steepness, conventionally characterized by a Hill coefficient *H* of the fitted Hill function, is limited by the number of TF binding sites *N* (Estrada et al., 2016). In this model, the promoter states *v*_*i*_, each associated with a transcription rate, are updated with every TF binding and unbinding event. When the model satisfies detailed balance, the forward and reversed transitions between any two states *v*_*i*_ and *v*_*j*_ are equilibrated:

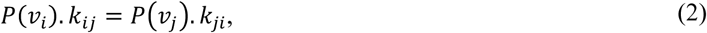

with *k*_*ij*_ and *k*_*ji*_ the forward and reverse rate constants, *P*(*v*_*i*_) and *P*(*v*_*j*_) the probability of the promoter states. Thus, the model can be collapsed into the linearized model as in Figure 4B, where only the number of occupied Bcd binding sites and not their identity matters. In this case, *H* cannot be above *N*. Experimental *H* values obtained when observing the protein level (Gregor, Tank, et al., 2007) and several features of the *hb* transcription dynamics (e.g. total amount of RNA produced (Garcia et al., 2013; Lucas, Tran et al., 2018)) range from ∼5 to ∼7, roughly equal to the number of known Bcd binding sites on *hb* promoter. This leads to assumptions that the 6 binding sites, with an unstable first Bcd-bound state (large *k*_−1_ in Figure 4B) and a stable fully bound state (small *k*_−*N*_ in Figure 4B), are sufficient to explain the observed pattern steepness in static measurements.

**Figure 4:**
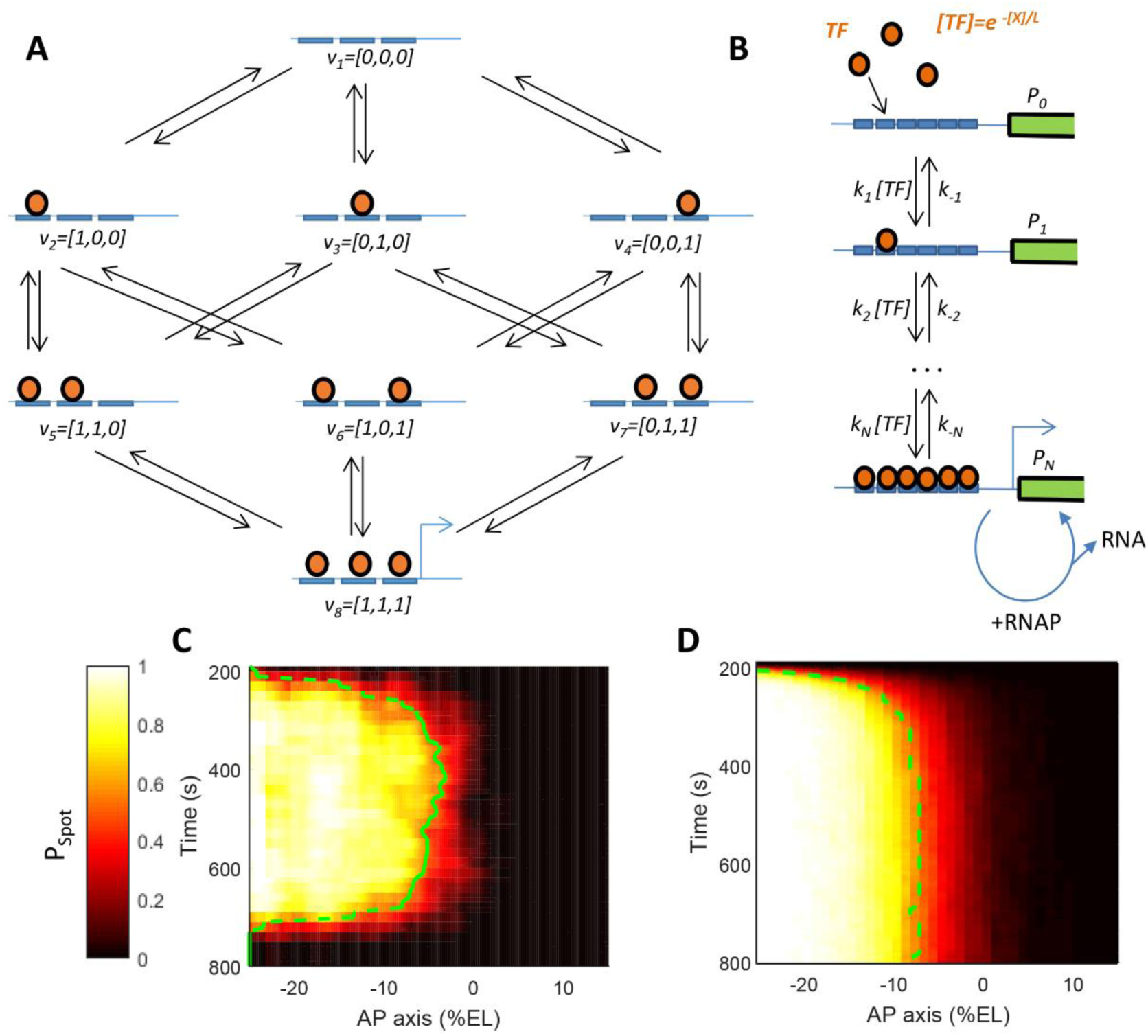
Modeling transcriptional regulation by the Bcd transcription factor through interactions with binding sites on *hb* promoter. **A)** A general model of transcription factors (TF) (orange) binding/unbinding to *N* binding sites of target promoter (*N*=3). Each node *v* corresponds to a unique ordered TF-bound promoter state. Figure adapted from (Estrada, Wong, DePace, & Gunawardena, 2016). **B)** A simplified promoter binding model assuming detailed balance to account for energy expenditure in the unbinding process (Estrada et al., 2016), coupled with transcription initiation. When many Bcd binding sites on the *hb* promoter are occupied, RNAP can randomly bind to the promoter and initiate transcription. **C-D)** The probability of an active transcription locus (P_Spot_, color bar) for the *hb* locus as a function of time in the nuclear cycle and position along the AP axis obtained from the MS2-MCP data (C, nc13) and from the model in B (D, with *N*=6, not accounting for mitosis at the end of interphase). Figure B-D are reused from (Lucas et al., 2018).

However, Estrada *et al.* did not consider the search time issue (Estrada et al., 2016) and their theory cannot explain how the *hb* pattern can be established in such a short time of 3 minutes following mitosis, as observed in live imaging data (Lucas, Tran et al., 2018) (Figure 4C). Considering a model which accounts for the Bcd search time for the *hb* promoter, Tran *et al* (Tran et al., 2018) found that, at the mid-boundary position, very high pattern steepness (*H* ≈ *N*) requires a very slow promoter switching time (called *τ*_*active*_) between Bcd-bound states allowing transcription and Bcd-free states prohibiting transcription. Therefore, according to the model, it should take a very long time for the pattern to be established and this is in contradiction with the experimental data from the MS2 system (Lucas, Tran et al., 2018). In addition, slow promoter dynamics results in high nuclei-to-nuclei variability in the amount of total RNA produced in each nuclear cycle. Thus, it would require even more averaging time to achieve the robust protein pattern (10% variability) observed in nuclear cycle 14 (Gregor, Tank, et al., 2007).

The failure of the simple model to explain both the observed high pattern steepness and fast establishment time begs for the reconsideration of the model’s assumptions. Most obvious candidates are either the underestimation of the number of Bicoid binding sites *N* or overestimation of the time it takes for Bcd to find the promoter *τ*_*search*_. Estrada *et al.* suggested that energy expenditure, which removes the detailed balance assumption from the binding and unbinding process, can expand the model’s limit on pattern steepness *H* beyond the binding site number *N*, allowing both high steepness and fast formation time at the same time (Estrada et al., 2016). However, including non-equilibrium binding does not resolve the problem of obtaining a steep yet precise boundary in a short time. Alternatively, Desponds *et al.* (Desponds, Vergassola, & Walczak, 2019) suggested that instead of probing the concentration for a fixed amount of time and then making the decision about the positioning of the nucleus, constantly updating the odds of being in an anterior *vs* posterior position always results in much faster decisions for a fixed accuracy. Assuming a promoter with 6 Bcd binding sites, they showed that the decision time can be reduced by an order of magnitude compared to the classical Berg-Purcell scheme, possibly below the 3 minute limit. Unlike the classical time averaging (Berg-Purcell) scheme, which theoretically can reach 100% accuracy given enough sensing time, the target gene “commits” to expression or silence when the odds favoring the anterior or posterior reach an acceptable confidence, even before the end of the exposure window. Possible molecular mechanisms for how this decision scheme could be implemented in fly embryos are still needed.

### 3.4. Dissecting noise in *hb* transcription

Noise in transcription dictates the variability of transcript and protein readouts after each interphase and might play a role in determining nuclei identity in downstream processes (Holloway et al., 2011). However, beyond its characterization from observed data, we still lack the mechanistic understanding of processes responsible for this noise.

As transcription bursts are prevalent across the embryo in the very short early nuclear cycles, *hb* transcription dynamics is well-fitted by a two-state model, in which the switching rates between the ON and OFF states are modulated by the nuclei’s position or Bcd concentration (Desponds et al., 2016; H. Xu et al., 2015; H. Xu, Skinner, Sokac, & Golding, 2016; Zoller et al., 2018). It should be noted that these ON and OFF states do not correspond to Bcd-free and Bcd-bound states of the promoter as in (Estrada et al., 2016; Tran et al., 2018): in the Bcd-saturating anterior region, the *hb* promoter is constantly active (i.e. bound by Bcd molecules) but transcription still occurs in bursts with the switching time between ON and OFF states ∼50*s* (Desponds et al., 2016). This suggests that promoter bursting may be an inherent property of transcription in this phase of development (Bothma et al., 2014; Zoller et al., 2018). The early transcription of *hb* is also regulated by other TFs such as maternal Hb (Lopes, Spirov, & Bisch, 2011; Porcher et al., 2010; Simpson-Brose et al., 1994) or Zelda (Harrison, Li, Kaplan, Botchan, & Eisen, 2011; Lucas et al., 2018; Nien et al., 2011; Z. Xu et al., 2014). Optogenetics was also used to inactivate the transcription factor Zelda in early embryos and reveals that this pioneer factor continuously regulates zygotic gene expression from nc10 to nc14 (McDaniel et al., 2019). Though these factors other than Bcd may not act as a source of positional information, their concentration may be rate-limiting and therefore responsible for bursts.

In the *hb* boundary region, where cell fate decision is critical, *hb* transcription readout is more variable than in the anterior region (Desponds et al., 2016; Lucas et al., 2018). This was initially thought to be due to extrinsic noise from Bcd variability (Gregor, Wieschaus, et al., 2007) being amplified in this region. However, given the very high steepness observed from the *hb* pattern (Lucas, Tran et al., 2018; H. Xu et al., 2015), the switching time between Bcd-dependent active and inactive states of the promoter (*τ*_*active*_) is expected to be at least one order of magnitude greater than the Bcd search time *τ*_*search*_ (Tran et al., 2018). If the search is done via 3D diffusion (*τ*_*search*_∼10*s*), *τ*_*active*_ at *hb* boundary is at least of a similar time scale as the switching time between ON and OFF states at the anterior (∼50*s*). In the context of very rapid embryo development (interphase duration of 5 to 15 minutes in nc11 to nc13), Bcd-dependent promoter switching becomes a non-negligible source of intrinsic noise that contributes substantially to the higher readout variability observed.

## 4. Perspectives

Despite several decades since the identification of the Bcd gradient, we still lack a quantitative description of the process allowing for the transcription of its main target gene *hb* in a step-like pattern. The short timescales of early development and its remarkable precision are questioning how fundamental limits coming from stochastic processes such as diffusion or bursty regulation influence the molecular encoding of regulation. To pursue these issues, recent experimental advances are allowing us to rigorously test theoretical ideas, and call for the creation of new models.

This simple example of developmental biology, is turning out to also be an ideal *in vivo* testbed for advances in single molecule techniques to study protein motility (Drocco et al., 2011; Mir et al., 2018) and promises to bring a more definitive view on how TF can find their target promoter and activate transcription. The different motilities of TF (Abu-Arish et al., 2010; Mir et al., 2018) and their inhomogeneous distribution in those so-called hubs (Mir et al., 2017) as seen with the Bcd case support the idea that the search time can be improved by 1D-diffusion along the DNA or via micro-environments of enriched TF concentration. However, with fluorescent-tagged molecules alone, it remains difficult to identify *in vivo* the location of the binding molecule along the chromosome, with enough resolution to detect 1D-diffusion or local enrichment. Conversely, such epigenomics tools as ChIP-seq give information on TF binding along the chromosome, but lack the temporal dynamics of these interactions. With advances in sample preparation and super-resolution microscopy, it might be possible to combine these two types of experiments to understand how TF searches for their specific sites. Another important question is the mechanisms of TF binding “cooperativity” after the search. In the Bcd case, binding cooperativity has been suggested in theoretical work to explain *hb*’s high pattern steepness at steady state (Estrada et al., 2016; Gregor, Tank, et al., 2007; Lopes et al., 2011; Tran et al., 2018). While high steepness is observed experimentally (corresponding to a Hill coefficient ∼7 to 8) (Lucas et al., 2018), *in vivo* measurements of Bcd binding cooperativity of such high order at the gene loci remains a challenge (Mir et al., 2018; H. Xu et al., 2015). *In vitro* studies showed that Bcd binding cooperativity is of low order (Hill coefficient ∼2 to 3) and robust to different target sites’ spacing (Ma et al., 1996), suggesting that it may originate from protein-protein interactions rather than from transcription factor-induced binding site conformational changes (Bray & Duke, 2004). More direct experimental evidence is required to fully elucidate high-order cooperativity observed in the expression pattern, possibly coming from different transcription factors.

Advances in live imaging of transcription offered a direct view on *hb* transcription pattern establishment in early nuclear cycles: the pattern is established in a considerable portion of the very short interphase and, in contrast to the low variability observed at the protein level, transcription is highly noisy, with prevalent bursty behavior across the embryo. This marks a shift of our approaches towards the understanding of transcription regulation during development from previously static and usually steady-state views to more theoretical and quantitative studies focusing on the out-of-steady-state dynamics of gene expression (Petkova et al., 2019; Tran et al., 2018). The question of TF searching for its targets remains of great interests to explain the rapidity of the *hb* pattern establishment despite a noisy transcription process. Transcription activation following TF binding, even though intensively studied for decades, also remains a mystery. In the context of development, it will be important to understand how the measurement process of positional information manages to combine the rapidity of the response with accuracy (Tran et al., 2018). These observations lead to suggestions that transcription activation may be non-reversible or involve transcriptional memory or mitotic bookmarking (Desponds et al., 2019; Ferraro, Esposito, et al., 2016; R. Zhao, Nakamura, Fu, Lazar, & Spector, 2011). The development of powerful optogenetic tools with high spatial and temporal resolution might shed light on this mechanism, as it is now possible to manipulate TF properties and observe the effects of these manipulations in real time under the microscope (Huang et al., 2017; McDaniel et al., 2019).

Finally, *hb* protein expression in nc14 is still remarkably robust, suggesting that temporal and spatial averaging downstream of noisy transcription is at work. Recent works have shown that positional information can be accurately decoded at the level of the gap genes (Petkova et al., 2019). However, decoding as well as encoding mechanisms in the earlier cell cycles remain unknown and the current experimental and theoretical methods are ready to tackle these questions. At last, it is likely that Bcd may no longer play a major role in maintaining *hb* pattern during the very long nuclear cycle 14 (Durrieu et al., 2018; Perry, Boettiger, & Levine, 2011; Perry et al., 2012). Given this, an intriguing aspect of this system is why it has been selected to provide such fast step-like pattern dynamics in the earlier nuclear cycles.

## Acknowledgements

We thank M. Andrieu, M. Coppey, G. Fernandes, C. Fradin, C. Perez-Romero, A. Ramaekers for stimulating discussions. Work in the Walczak and Dostatni labs is supported by PSL IDEX REFLEX Grant for Mesoscopic Biology (AMW & ND), ANR-11-BSV2-0024 Axomorph (AMW & ND), ARC PJA20151203341 (ND) and ANR-11-LABX-0044 DEEP Labex (ND). HT was supported by the PSL IDEX REFLEX Grant, the Institut Curie and the DEEP Labex.

